# The inhibitory activity of glycine transporter 2 targeting bioactive lipid analgesics are influenced by formation of a deep lipid cavity

**DOI:** 10.1101/2020.09.10.290908

**Authors:** Katie A. Wilson, Shannon N. Mostyn, Zachary J. Frangos, Susan Shimmon, Tristan Rawling, Robert J. Vandenberg, Megan L. O’Mara

## Abstract

The human glycine transporter 2 (GlyT2 or SLC6A5) has emerged as a promising drug target for the development of new analgesics to manage chronic pain. N-acyl amino acids inhibit GlyT2 through binding to an allosteric binding site to produce analgesia in vivo with minimal overt side effects. In this paper we use a combination of medicinal chemistry, electrophysiology, and computational modelling to explore the molecular basis of GlyT2 inhibition at the allosteric site. We show how N-acyl amino acid head group stereochemistry, tail length and double bond position promote enhanced inhibition by deep penetration into the binding pocket. This work provides new insights into the interaction of lipids with transport proteins and will aid in future rational design of novel GlyT2 inhibitors.

## Introduction

One in ten adults worldwide are diagnosed with chronic pain each year (1). Despite the high rate of chronic pain there is a lack of safe and effective treatment options, which in turn has significant social and economic consequence (2). In the mammalian central nervous system, the neurotransmitter glycine inhibits the pain signalling pathway (3). Synaptic concentrations of glycine are controlled by the two glycine transporters, GlyT1 and GlyT2, which are responsible for clearing glycine from synapses (4). GlyT2 is expressed by presynaptic neurons and is also responsible for replenishing presynaptic glycine concentrations to maintain glycinergic neurotransmission. Inhibitors of GlyT2 slow the reuptake of glycine to prolong glycine neurotransmission (5) and have been developed as potential therapeutics in the treatment of chronic pain (6–11).

N-acyl amino acids comprised of an amino acid head group conjugated via an amide bond to a monounsaturated lipid tail represent a novel class of GlyT2 inhibitors (6). One of the most promising bioactive lipids from this series, oleoyl D-lysine (C18ω9-D-Lys), is a selective and potent GlyT2 inhibitor that is metabolically stable, blood brain barrier permeable, and produces analgesia in a rat model of neuropathic pain with minimal side effects (12). The reduced toxicity of these compounds is attributed to partial inhibition of GlyT2. They reduce, but do not completely block glycine transport, which allows presynaptic glycine concentrations to be maintained for subsequent repackaging into synaptic vesicles to maintain glycinergic neurotransmission (13). The most potent bioactive lipids bear positively charged (Lys) or aromatic (Trp) amino acid head groups and inhibit GlyT2 with IC_50_ concentrations of less than 50 nM (12). Within the allosteric binding site, oleoyl-L-lysine (C18ω9-L-Lys) orients tail-down so that the tail intercalates between aliphatic-rich regions of TM5 and TM8 and the oleoyl double bond is in close proximity to TM5 (6). The amino acid head group is accessible to the extracellular solution and stabilised by aromatic residues in TM7, TM8, and EL4 (6).

In the present work we use a combination of medicinal chemistry, electrophysiology, and computational modelling to explore the structure-activity relationship for lipid inhibitors of GlyT2 to understand how head group stereochemistry and chemical features of the acyl tail affect inhibitor activity and interaction with GlyT2 at a molecular level. The effects of head group stereochemistry are investigated for the most potent previously identified lipid inhibitors, C18ω9-Lys and C18ω9-Trp. To further understand the effect of structure on inhibitor potency, a series of acyl-lysine analogues were synthesised with variations in the tail length (C18, C16 and C14) and position of the double bond within the lipid tail (ω9, ω7, ω5 and ω3; Figure 1). Molecular dynamics (MD) simulations were used to provide a structural basis for the effect of chemical changes on lipid inhibitor properties. We demonstrate that the potency of the lipid inhibitors was greatly influenced by a combination of both the stereochemistry of the head group, and the length and saturation of the lipid tail. Furthermore, inhibitor potency is shown to depend on deep penetration of the lipid tail into a stabilized location in the allosteric binding site. This study develops a comprehensive structure-activity relationship for lipid inhibitors of GlyT2. This is critical for future rational design of more effective GlyT2 inhibitors for the treatment of chronic pain and may have broader implications for modulation of other SLC6 transporters.

**Figure 1.**
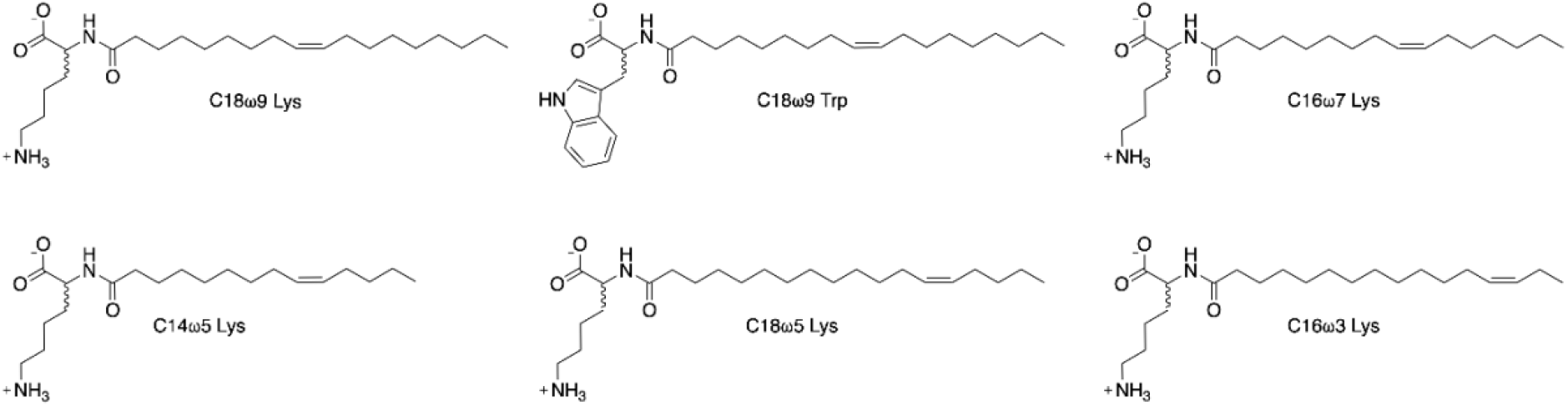
Chemical structures of GlyT2 lipid inhibitors considered in the present work. All inhibitors were synthesised as enantiopure D- and L-isomers.

## Methods

All N-acyl amino acids were synthesised as previously described (Mostyn et al., 2019). 10 mg mL-1 stock solutions of N-acyl amino acids were dissolved in DMSO and diluted in frog ringers solution (96 mM NaCl, 2 mM KCl, 1 mM MgCl2, 1.8 mM CaCl2, 5 mM HEPES, pH 7.5) to desired concentrations. Final solutions contained 0.0025% DMSO, a concentration which had no effect on transporter or receptor function.

### Wild type (WT) and mutant RNA transcription

Human GlyT2a WT cDNA was subcloned into the plasmid oocyte transcription vector (pOTV). The amplified GlyT2/pOTV product was then transformed in E. coli cells, and subsequently purified using the PureLink Quick Plasmid Miniprep Kit (Invitrogen by Life Technologies, Löhne, Germany), and sequenced by the Australian Genome Research Facility (Sydney, Australia). The purified plasmid DNA was linearised via the restriction enzyme, SpeI (New England Biolabs (Genesearch), Arundel, Australia) for GlyT2a. Complementary RNAs were synthesised using the mMESAGE mMACHINE T7 kit (Ambion, Texas, USA).

### Oocyte preparation and injection

All work involving the use of animals was performed in accordance with the Australian Code of Practice for the Care and Use of Animals for Scientific Purposes. Xenopus laevis frogs (NASCO, Wisconsin, USA) were anesthetised with 0.17% (w/v) 3-aminobenzoic acid ethyl ester and had an ovarian lobe removed via an incision in the abdomen. Stage V oocytes were isolated from the lobe via digestion with 2 mg mL-1 collagenase A (Boehringer, Mannheim, Germany) at 26°C for 1 hour. 20 ng of cRNA encoding GlyT2 was injected into each oocyte cytoplasm (Drummond Nanoinject, Drummond Scientific Co., Broomall, Pennsylvania, USA). The oocytes were then stored in frog Ringer’s solution (96 mM NaCl, 2 mM KCl, 1 mM MgCl2, 1.8 mM CaCl2, 5 mM HEPES, pH 7.5) which was supplemented with 2.5 mM sodium pyruvate, 0.5 mM theophylline, 50 μg/mL gentamicin and 100 μM mL-1 tetracycline. The oocytes were stored at 18°C for 3-5 days, until transporter expression was adequate for measurement using the two-electrode voltage clamp technique.

### Two-electrode voltage clamp electrophysiology

GlyT2 is electrogenic, allowing activation to be measured via the two-electrode voltage clamp technique. Oocytes were voltage clamped at −60 mV, and whole-cell currents generated by the substrate were recorded with a Geneclamp 500 amplifier (Axon Instruments, Foster City, California, USA), digitised by a Powerlab 2/20 chart recorder (ADInstruments, Sydney, Australia). LabChart version 8 software (ADInstruments, Axon Instruments) was used to visualise and process current traces. Recordings were performed in frog Ringer’s solution. Oocytes were placed in an oval-shaped bath with a volume of 0.3 mL, with laminar flow around the oocyte at a rate of 12 mL min-1 under gravity feed.

Each of the synthesised acyl-lysine compounds were first tested for their inhibitory activity by coapplying the analogues with 30 μM glycine and assessing their ability to reduce glycine transport currents. Increasing concentrations of each acyl-lysine inhibitor was applied to generate concentration response curves, from which IC_50_ values and percent max inhibition values were calculated. Acyl-lysines were also applied to the related transporter GlyT1 to assess their selectivity and were shown to not significantly inhibit GlyT1 currents at concentrations up to 3 μM. We previously showed that concentrations of up to 3 μM of oleoyl L-lysine do not alter uninjected oocytes, and that the critical micelle concentration of positively charged bioactive lipids is above this value.

### Computational Modeling

Docking and molecular dynamics simulations were performed using the protocol developed in our previous work (6). Briefly, each of the lipid inhibitors was docked to our previously validated GlyT2 homology model using Autodock vina (14). The lipids were treated in a united atom representation to be consistent with the subsequent MD simulations and treated as flexible. Lipid inhibitors were docked to the previously identified extracellular allosteric binding pocket. Following docking, the pose with the lipid inhibitors bound with the tail inserted into the extracellular pocket and the double-bond in close proximity to TM5 and the protein-lipid interface, was considered for simulation.

The Automated Topology Builder and Repository (ATB) (15) was used to develop united atom coordinates and parameters for the lipid inhibitors (Molecule IDs: 252930, 340331, 252919, 253354, 364924, 364925, 364971, and 364972). To ensure that there was no isomerisation around the *cis* double bond, the force constant related to this dihedral angle in the lipid inhibitors was adjusted from 5.86 kJ mol^−1^ rad^−2^ to 41.80 kJ mol^−1^ rad^−2^. The protonation state for all lipids was that in which it would most likely be found at physiological pH (pH 7): POPC and the lipid inhibitors with a Lys head group were zwitterions, while the lipid inhibitors with a Trp head group were deprotonated. The simulations were performed using GROMACS version 2016.1 (16), with the GROMOS 54A7 force field for lipids and proteins (17). GlyT2 was embedding in a membrane containing 80% POPC and 20% CHOL, each system was neutralized, and salt was added to a concentration of 0.15 M NaCl. All systems were minimized using a steepest descent and equilibrated with decreasing restraints on the protein in five sequential 1 ns simulations (1000 kJ mol^−1^ nm^−2^, 500 kJ mol^−1^ nm^−2^, 100 kJ mol^−1^ nm^−2^, 50 kJ mol^−1^ nm^−2^, and 10 kJ mol^−1^ nm^−2^. Each system was then simulated without constrains in triplicate for 500 ns. Throughout all simulations the pressure was maintained at 1 bar using semi-isotropic position scaling and the Berendsen barostat (τP = 0.5 ps and isothermal compressibility = 4.5×10^−5^ bar), the temperature was maintained at 300 K using the Bussi-Donadio-Parrinello velocity rescale thermostat (τP = 0.1 ps), the periodic boundary condition, a 2 fs timestep, SETTLE was used to constrain the geometry of water molecules and LINCS was used to constrain the covalent bond lengths of the solute. Visualization of the simulations was performed using the Visual Molecular Dynamics software (VMD) (18). Analysis was done on frames spaced by 0.1 ns using gromacs tools, and VMD.

## Results

### Head group stereochemistry affects C18ω9-Trp binding but not C18ω9-Lys binding

Bioactive lipids bearing Lys or Trp head groups in the L-configurations are among the most potent GlyT2 inhibitors in the series, inhibiting GlyT2 with IC_50_ concentrations of 25.5 nM and 54.6 nM, respectively (Table 1). Interestingly, when the head group is converted to the D-configuration the Lys analogue (C18ω9-D-Lys) retains potency while the Trp analogue (C18ω9-D-Trp) is inactive (Table 1) (6). To investigate the molecular basis of this activity, 500 ns MD simulations were performed in triplicate, in which one molecule of either the L- or D-stereoisomers of C18ω9-Lys was docked into the extracellular allosteric binding site of our GlyT2 homology model (19). Only binding poses in which the lipid tail was inserted into the extracellular allosteric binding site and the double-bond was in close proximity to TM5 were considered (6). Regardless of head group amino acid type or stereochemistry, the transporter remains in an outward occluded conformation (Table S1) and the membrane properties are not affected by lipid inhibitor binding (Table S2).

**Table 1.** Activity of acyl-lysine analogues at GlyT2 and GlyT1

|  | L-Headgroup Stereochemistry |  |  | D-Headgroup Stereochemistry |  |  |
| --- | --- | --- | --- | --- | --- | --- |
|  | IC<sub>50</sub> (nM) | % max inhibition | Activity at GlyT1 | IC<sub>50</sub> (nM) | % max inhibition | Activity at GlyT1 |
| C18 ω9 Trp | 54.6<sup>a</sup> | 86.2<sup>a</sup> | >10 μM<sup>a</sup> | >10 μM<sup>a</sup> | 14.2<sup>a</sup> | >10 μM<sup>a</sup> |
| C18 ω9 Lys | 25.5<sup>a</sup> | 86.8<sup>a</sup> | >3 μM<sup>a</sup> | 48.3<sup>a</sup> | 91.0<sup>a</sup> | >3 μM<sup>a</sup> |
| C16 ω7 Lys | 66.6<br>(49.9 – 88.7) | 90.2 ± 2.1 | TBC | 602<br>(373 – 856) | 96.2 ± 2.1 | >3 μM |
| C14 ω5 Lys | 703<br>(414 – 1250) | 96.2 ± 7.3 | TBC | 1380<sup>b</sup> | 79.7 ± 3.6 | >3 μM |
| C18 ω5 Lys | 67.5<br>(31.7 – 143) | 81.3 ± 5.1 | TBC | 64.9<br>(36.3 – 123) | 87.6 ± 3.7 | >3 μM |
| C16 ω3 Lys | 10.8<br>(8.37 – 13.8) | 94.9 ± 1.6 | TBC | 699<br>(343 – 1480) | 91.8 ± 8.6 | >3 μM |

<sup>a</sup>previously published data from Mostyn et al., 2019a, <sup>b</sup>95% CI was not able to be calculated.

Both C18ω9-L-Lys and C18ω9-D-Lys remain bound in the allosteric site throughout the combined 1500 ns of MD simulation with the lipid tail in an extended conformation (~17-20 Å measured from the end of the tail to the stereocenter, Figure S1), intercalated between TM5, TM7, and TM8 (Figure 2a and Figure S2). The amino acid head groups remains in close proximity to the protein/bilayer-water interface and interacts with the extracellular regions of TM5, TM7, TM8, and EL4, including hydrogen bonding with cationic arginine residues (i.e., R439, R531 and R556) and stacking with non-polar residues (i.e., F526 and W563). The tail is positioned in a hydrophobic pocket lined by L436, V523, Y550, A553, F567 (Figure 2b), in agreement with previous studies (6). Interactions that occur with both the head group and tail are sustained for >75% of the total simulation time (Table S3). Notably, the more potent inhibitor, C18ω9-L-Lys, has deeper penetration into the binding pocket than C18ω9-D-Lys. Y550 is a key residue in the interaction of both isomers. Y550 interacts with C18ω9-L-Lys above the oleoyl double bond. The change of stereochemistry to C18ω9-D-Lys decreases the depth of penetration of the oleoyl tail and shifts the interaction with Y550 below the double bond of C18ω9-D-Lys. In this position, Y550 hydroxyl forms hydrogen bonds with the backbone of W563, so that the lipid tail is sandwiched between Y550 and W563, with W563 interacting directly with the double bond with a CH-π interaction. The interactions of Y550 and W563 with both isomers correlates with mutagenesis data reporting the GlyT2 Y550L and W563L mutants are not inhibited by C18ω9-L-Lys or C18ω9-D-Lys (6).

**Figure 2.**
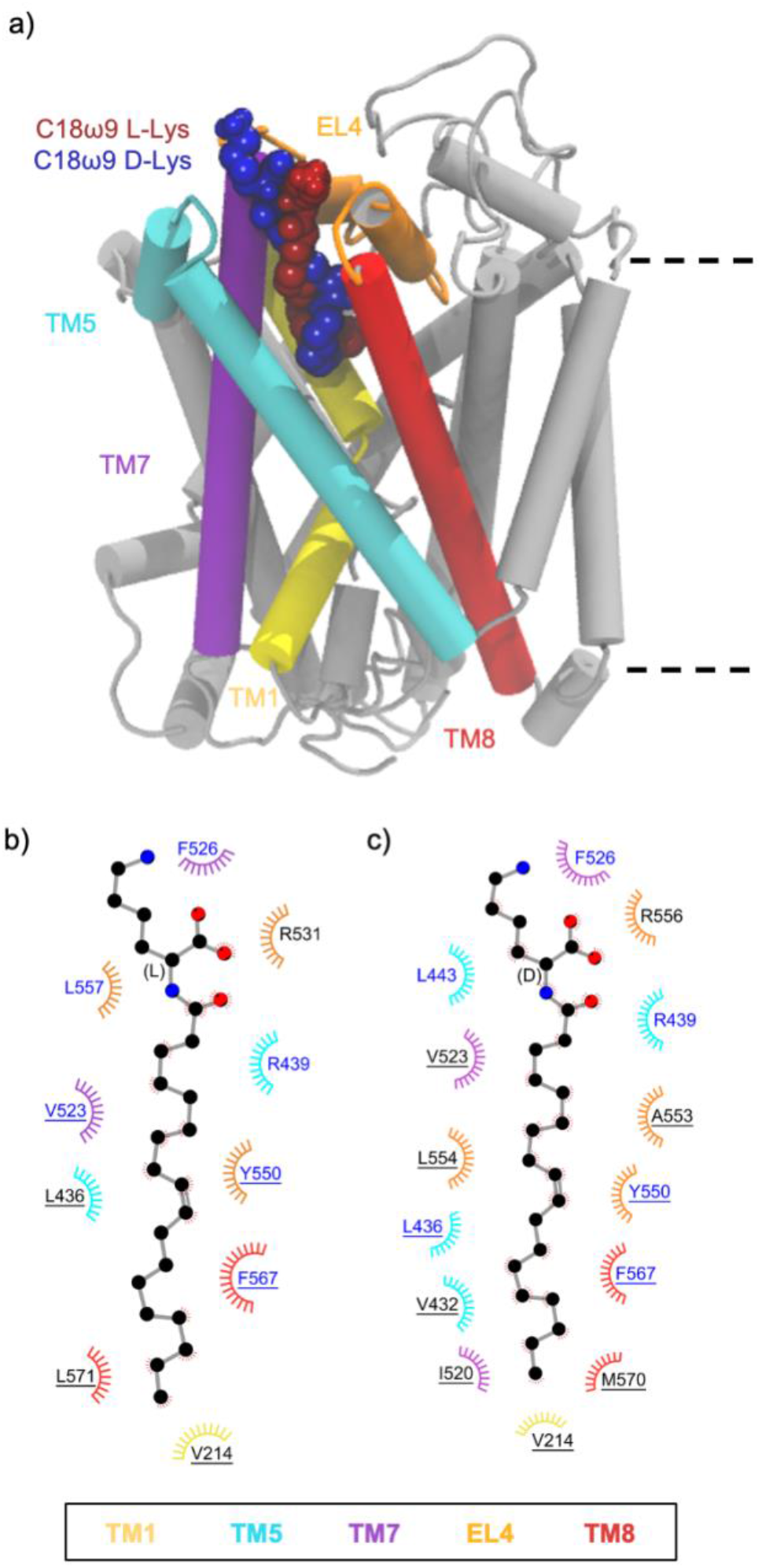
a) Positions of the Lys C18ω9 lipid inhibitor binding in the extracellular allosteric binding pocket of GlyT2. b) and c) Residues that interact with b) C18ω9-L-Lys or c) C18ω9-D-Lys in representative snapshots from MD simulations of the lipid inhibitors bound in the allosteric binding site. Residues that interact with the lipid inhibitor for >75% or <75% of the total simulation time are highlighted in blue or black, respectively. Residues are split based on whether the amino acid interacts with the lipid inhibitor head-group (no underline) or tail-group (underlined).

In contrast to the lysine-based analogues, the potency of the bioactive lipids bearing Trp head groups is significantly impacted by stereochemistry such that the D-isomer is inactive against GlyT2 (Table 1) (6). In MD simulations C18ω9-L-Trp bound to the GlyT2 allosteric site in a similar conformation as C18ω9-L-Lys. C18ω9-L-Trp is again intercalated between TM5, TM7, and TM8 with the double-bond in close proximity to TM5 (Figure 3a and Figure S3). The D-isomer, C18ω9-D-Trp, alters its orientation around the stereocenter so the C18ω9 tail points towards EL4, rather than facing the pocket formed by TM5, TM7 and TM8. While C18ω9-D-Trp still binds to the GlyT2 extracellular allosteric site, it adopts a much more curled conformation than C18ω9-L-Trp (~13 vs ~18 Å measured from the end of the tail to the stereocenter; Figure S1). This curled conformation results in a shallower binding interaction and reduced depth of penetration of C18ω9-D-Trp into the extracellular allosteric site which alters the coordination of the lipid tail within the binding pocket. Key head group interactions with R439 and F526 were maintained. Residues L436, V523, Y550, A553, and L557 that interact with the lipid tail of C18ω9-D-Lys, C18ω9-L-Lys and C18ω9-L-Trp instead form interactions with the head group of C18ω9-D-Trp (Figure 3b and Table S3). Only one interaction forms with the C18ω9-D-Trp lipid tail for >75% of the simulation time (Figure 3c and Table S3). Without these key residues stabilizing the position of the acyl tail of C18ω9-D-Trp within the binding site, the tail does not remain bound between TM5, TM7 and TM8 over the course of the MD simulation as observed for the active lipid inhibitors. Instead the lipid tail leaves the allosteric binding site and reorients in the solution towards EL4, where it adopts a variety of conformations. Despite C18ω9-D-Trp remaining bound in the extracellular allosteric binding site throughout the total simulation time and the seemingly favourable interaction of the C18ω9-D-Trp head group, no inhibition of GlyT2 is achieved. This indicates that in order for the bioactive lipids to inhibit GlyT2, the tail must be stabilized within the extracellular allosteric binding site, positioned between TM5, TM7 and TM8. This is in agreement with the inability of free amino acids to cause inhibition (20).

**Figure 3.**
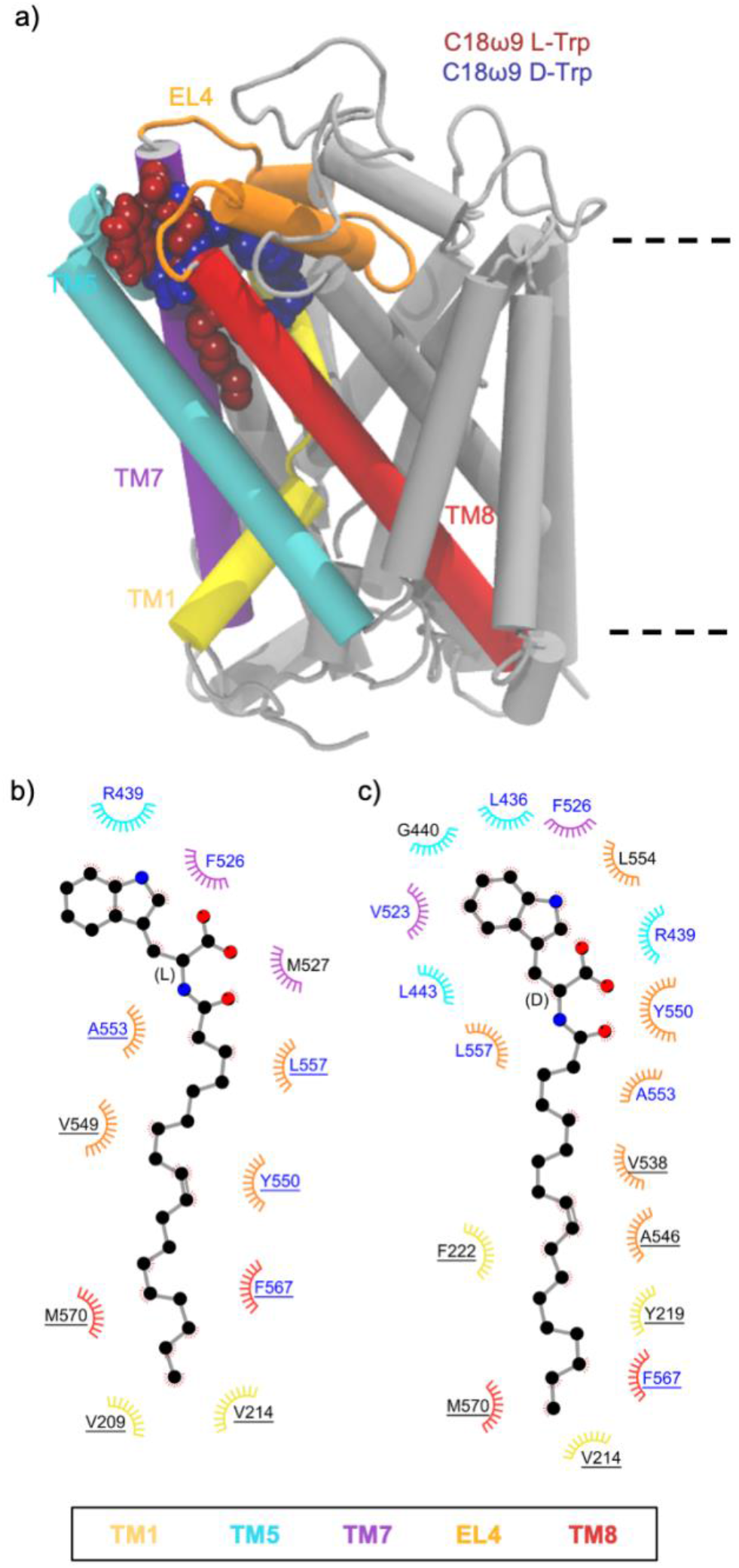
a) Position of the two Trp C18ω9 lipid inhibitors in the extracellular allosteric binding pocket of GlyT2. b) and c) Residues that interact with b) C18ω9-L-Trp or c) C18ω9-D-Trp in representative snapshots from MD simulations of the lipid inhibitors bound in the allosteric binding site. Residues that interact with the lipid inhibitor for >75% or<75% of the total simulation time are highlighted in blue or black, respectively. Residues are split based on whether the amino acid interacts with the lipid inhibitor head-group (no underline) or tail-group (underlined).

### Shortening the tail of acyl-lysine analogues reduces the depth of penetration and inhibitor potency

The lipid inhibitors described above contain 18 carbon acyl tails with a *cis*-double bond in the Δ9 position (i.e. 9 bonds from the amide linkage to the amino acid head group), and penetration of the lipid tails into the allosteric site appears to be a critical determinant of inhibitory activity. To investigate the effect of tail length on GlyT2 inhibitory activity we synthesised D- and L-lysine-based inhibitors with truncated tails. Double bonds were maintained in the Δ9 position and overall chain lengths was reduced to C16 or C14 (C16ω7-Lys and C14ω5-Lys, respectively). The chemical structures of these new lipids are shown in Figure 1 and their synthesis and characterisation is described in the Supplementary Material. Newly synthesised acyl lysine analogues were then tested against GlyT2 and also tested for selectivity by testing against the closely related GlyT1 transporter using two-electrode voltage clamp electrophysiology (see Table 1 for inhibitory data).

While C18ω9-L-Lys and C18ω9-D-Lys had similar levels of activity, head group conformation greatly affected the potencies of the chain shortened analogues. C16ω7-L-Lys inhibited GlyT2 with an IC_50_ of 66.6 nM, but the corresponding D-isomer was 9-fold less potent (IC_50_ of 602 nm, Figure 4b-c). Further shortening of the tails to Lys C14ω5 produced a similar preference for L-vs D-, albeit with markedly decreased the potency compared to the C18 analogues (IC_50_ concentrations of 770 and 1380 nM, for Lys-L-C14ω5 and Lys-D-C14ω5 respectively; Figure 4d).

**Figure 4.**
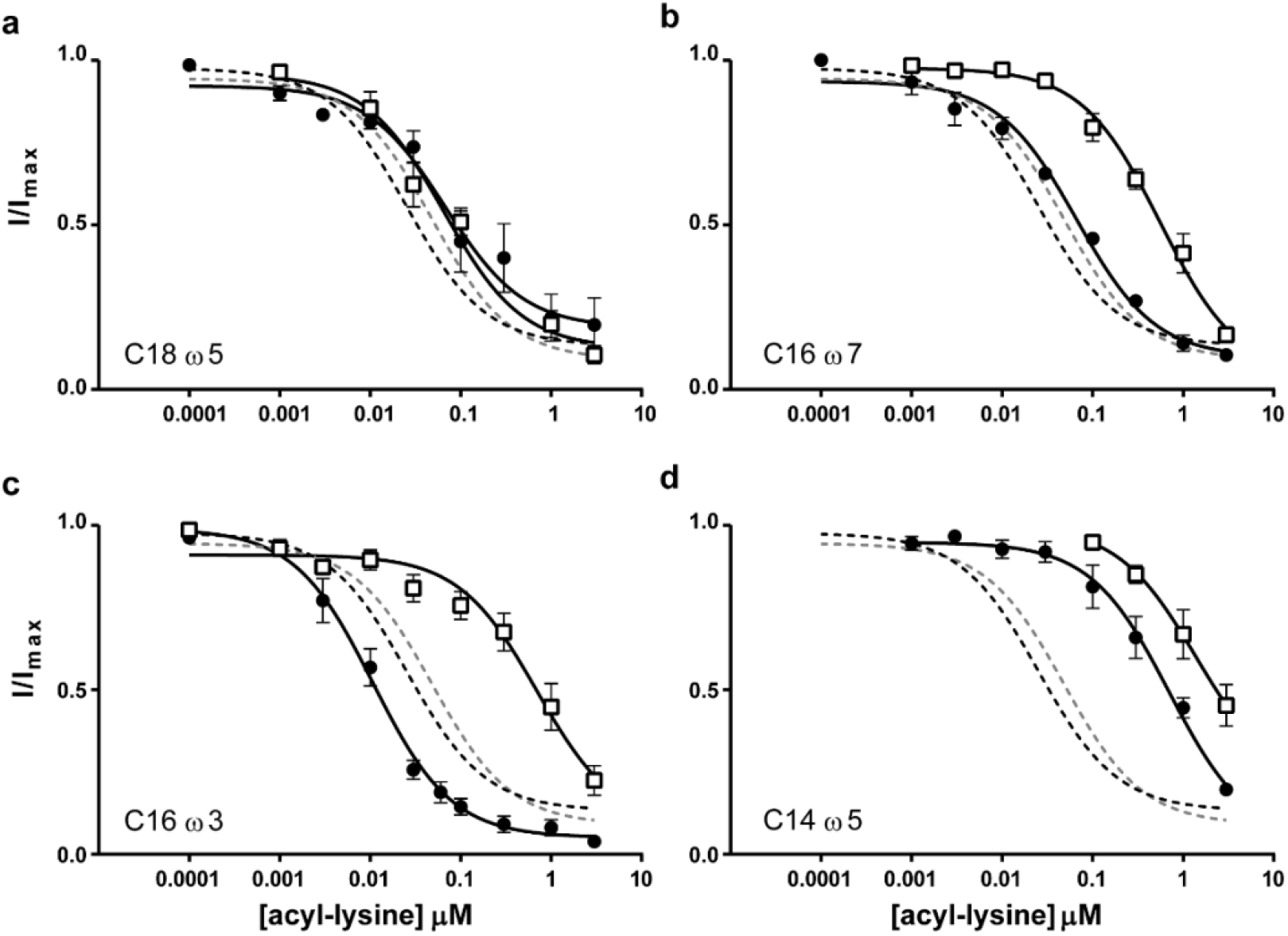
Inhibitory activity of acyl-lysine analogues at GlyT2. Increasing concentrations of acyl lysine compounds were applied to oocytes expressing wild type GlyT2 transporters. The transport current at each concentration is normalised to the current produced by 30 μM glycine in the absence of inhibitor. For each panel the L-isomer is represented by black circles and the D-isomer is shown as open squares. The curve fits for oleoyl (C18ω9) lysine are also shown as black (L-) and grey (D-) dashed lines for comparison. a) C18ω5 acyl-lysine. b) C16ω7 acyl-lysine, c) C16ω3 acyl-lysine d) C14ω5 acyl-lysine.

To provide a structural explanation of this difference in activity with changing tail length, C16ω7 and C14ω5 acyl lysines in the L- and D-configurations were docked to the extracellular allosteric binding site, and 500 ns of unrestrained MD simulations were performed in triplicate. Throughout all simulations, GlyT2 remains in an outward occluded conformation, regardless of the tail length of the acyl lysines (Table S1). The membrane properties are not affected (Table S2).

MD simulations showed both C16ω7-L-Lys and C14ω5-L-Lys remained bound in the allosteric binding site throughout the simulation, with the tail positioned between TM5, TM7 and TM8 (Figure 5a and Figure S4). The Lys head groups of both C16ω7-L-Lys and C14ω5-L-Lys remain in close proximity to the protein/bilayer-water interface, interacting with the extracellular regions of TM5, TM7, TM8, and EL4. C16ω7-L-Lys adopts a similar orientation to that observed for C18ω9-L-Lys, in close proximity to the key binding pocket residues (Figure 5b and Table S4). As was observed for C18ω9-L-Lys, the C16ω7-L-Lys head group interacts with R436 and F526. Similarly, the L-Lys C16ω7 tail is located between TM5, TM7 and TM8 in the extracellular allosteric binding site where it interacts with L436, V523, Y550, L557, and F567 and the bottom of the pocket is flanked by V214. As was the case for C18ω9-L-Lys, Y550 interacts with the lipid tail just above the double bond of C16ω7-L-Lys. The similarities between the overall orientation and interactions of C18ω9-L-Lys and C16ω7-L-Lys with key residues in the binding pocket provides a structural basis for the potent inhibition of GlyT2 by both molecules. Further shortening of the lipid tail to give C14ω5-L-Lys significantly reduces the depth of the tail penetration into the binding pocket (Figure 5a). Furthermore, shortening of the lipid tail dramatically alters C14ω5-L-Lys head group interactions. The C14ω5-L-Lys head group interacts with the membrane, forming hydrogen bonds with POPC head groups (Figure S5). There are no interactions with the C14ω5-L-Lys head group that persisted for >75% of the total simulation time (Figure 5c and Table S4). In contrast, the tail maintains interactions with key residues (i.e., L436, V523, Y550, and F567) for >75% of the total simulation time. In both C14ω5-L-Lys and C16ω7-L-Lys, the tail only penetrates as deep as F567 (Figure 5b and 5c), which interacts with C6 of C18ω9-L-Lys and C5 of C16ω7-L-Lys. This change in orientation of C14ω5-L-Lys (IC_50_ of 703 nm) relative to C16ω7-L-Lys (IC_50_ of 66.6 nm) and C18ω9-L-Lys (IC_50_ of 25.5 nm) correlates with the dramatic decrease in GlyT2 inhibition observed for C14ω5-L-Lys.

**Figure 5.**
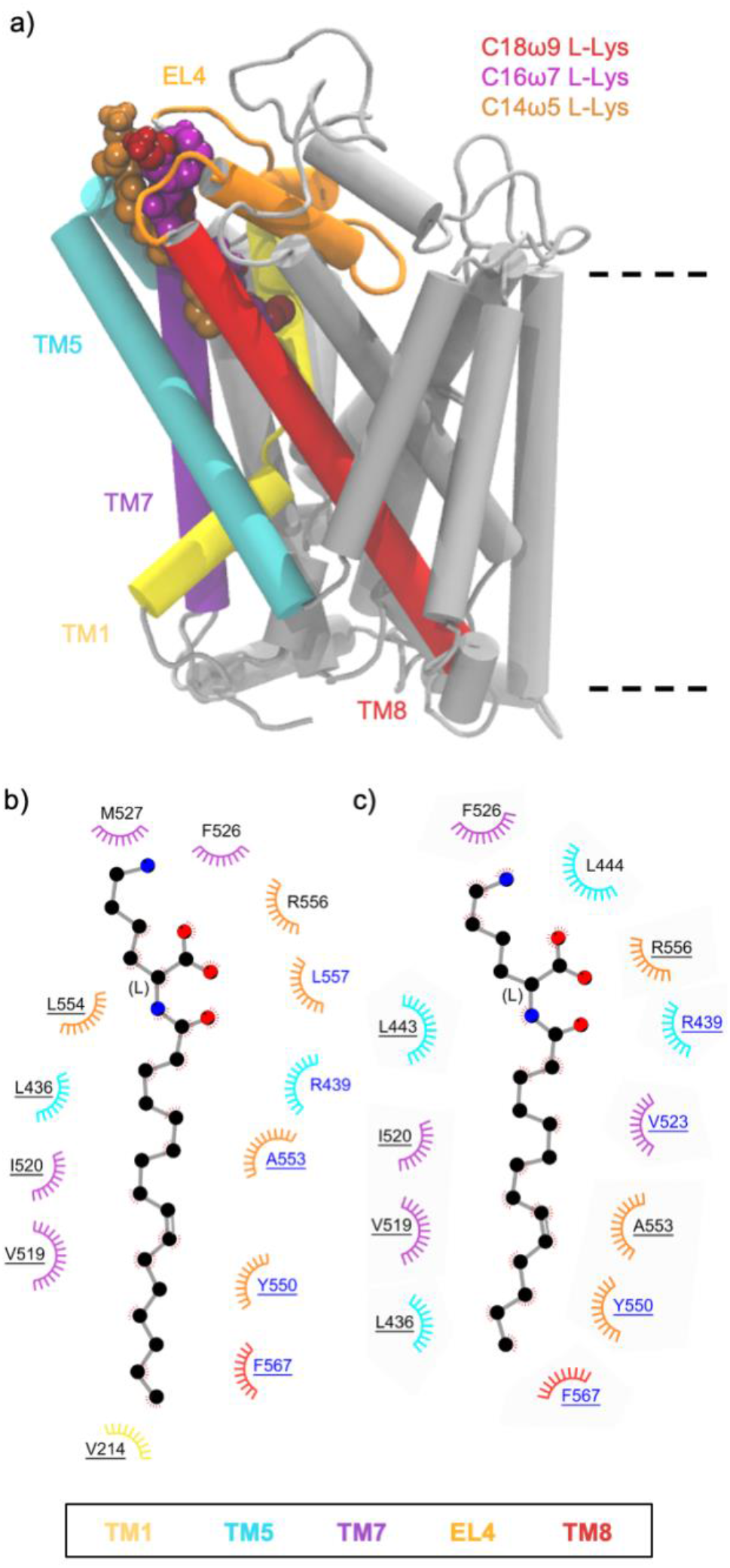
a) Positions of the L-Lys C18ω9, C16ω7 and C14ω5 lipid inhibitor binding in the extracellular allosteric binding pocket of GlyT2. b) and c) Residues that interact with C16ω7-L-Lys b) or C14ω5-L-Lys c) in representative snapshots from MD simulations of the lipid inhibitors bound in the allosteric binding site. Residues that interact with the lipid inhibitor for >75% or <75% of the total simulation time are highlighted in blue or black, respectively. Residues are split based on whether the amino acid interacts with the lipid inhibitor head-group (no underline) or tail-group (underlined).

On change of stereochemistry to the D-configuration, C16ω7-D-Lys and C14ω5-D-Lys only remain bound in the extracellular allosteric site for approximately one third (~500 ns) of the total simulation time. This is in contrast to C18ω9-D-Lys, which remains bound throughout the entire simulation. Furthermore C16ω7-D-Lys and C14ω5-D-Lys adopt a different conformation in the extracellular allosteric binding pocket to C18ω9-D-Lys (Figure 6a and Figure S6). Specifically, the C16ω7-D-Lys head group interacts with the head groups of membrane POPC lipids (Figure S4b) while maintaining interactions with key GlyT2 residues (F526, R439 and R556; Figure 6b and Table S4). While the C16ω7-D-Lys tail group is positioned in the extracellular allosteric binding pocket in a similar orientation to C18ω9-D-Lys and interacts with L436, V523, Y550, and F567, none of these interactions persist for >75% of the total simulation time (Figure 6b and Table S4). The reduced occupancy and different binding orientation of C16ω7-D-Lys (IC_50_ of 602 nm) in the extracellular allosteric binding site in part explain the 12-fold reduced potency relative to C18ω9-D-Lys (IC_50_ of 48.3 nm). In the case of C14ω5-D-Lys, the D-Lys head group interacts with F526 and R439 in a similar manner to C18ω9-D-Lys. However, the C14ω5-D-Lys tail protrudes into the surrounding membrane where it interacts with POPC at the protein-lipid interface (Figure S7). In this orientation, L443 is the only key residue interacting with the lipid tail for >75% of the total simulation time. The lack of interactions and reduced occupancy in the extracellular allosteric binding site may in part explain why C14ω5-D-Lys is not an effective inhibitor of GlyT2 (IC_50_ of 1380 nm).

**Figure 6.**
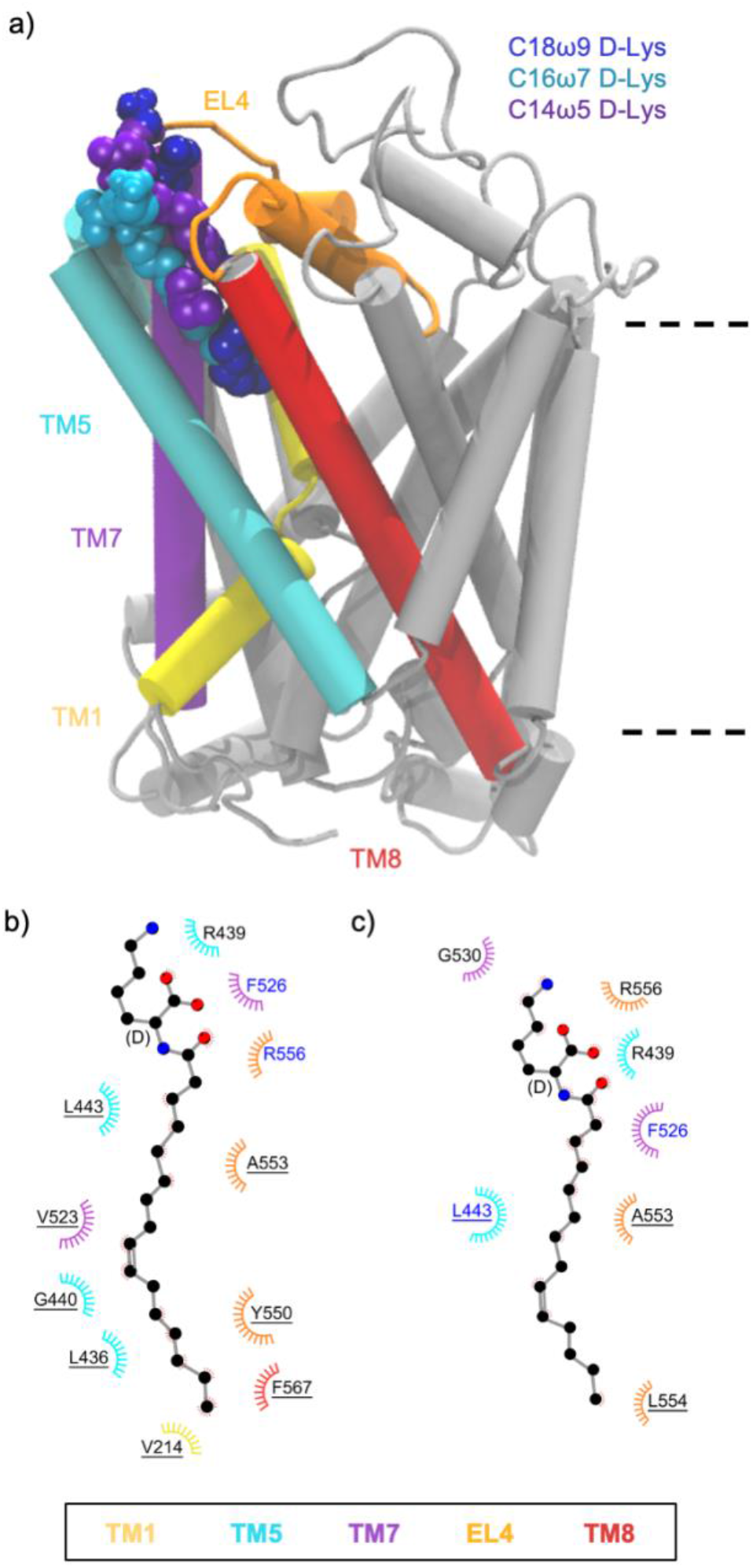
a) Positions of the D-Lys C18ω9, C16ω7 and C14ω5 lipid inhibitor binding in the extracellular allosteric binding pocket of GlyT2. b and c) Residues that interact with C16ω7-D-Lys (b) or C14ω5-D-Lys (c) in representative snapshots from MD simulations of the lipid inhibitors bound in the allosteric binding site. Residues that interact with the lipid inhibitor for >75% or <75% of the total simulation time are highlighted in blue or black, respectively. Residues are split based on whether the amino acid interacts with the lipid inhibitor head-group (no underline) or tail-group (underlined).

### The position of the double bond in the lipid tail changes the activity of acyl-lysine analogues

To assess the effect of the double bond position on activity we synthesised C18ω5-Lys and C16ω3-Lys, which contain *cis*-double bonds in the Δ13 position (Figure 1). Both L- and D-isomers of C18ω5-Lys inhibit GlyT2 with IC_50_ concentrations of 67.5 and 64.9 nM, respectively (Table 1 and Figure 4a). This trend is consistent with the results for C18ω9-L-Lys and C18ω9-D-Lys, where the configuration of the amino acid head group did not greatly alter the activity. However, when the length of the acyl chain was decreased to C16 the position of the double bond produced a marked difference in inhibitory activity between isomers. Thus, C16ω3-L-Lys is a potent inhibitor of GlyT2 (IC_50_ of 10.8 nM) while the corresponding D-isomer was 65-fold less potent, with an IC_50_ of 699 nm (Figure 4b-c).

To provide a structural explanation of this difference in activity with changing tail length, the Land D-isomers of C18ω5 and C16ω3 Lys were docked to the extracellular allosteric binding pocket and simulated for 500 ns in triplicate. Regardless of the presence of a bound inhibitor, GlyT2 again remains in an outward occluded conformation (Table S1) throughout the simulations and the membrane properties are not affected (Table S2). Both stereoisomers with C18 tails (C18ω5-L-Lys and C18ω5-D-Lys) remained bound in the allosteric pocket throughout all simulations. The overall binding conformation and potencies of C18ω5-L-Lys and C18ω5-D-Lys were similar to that of the C18ω9-L-Lys and C18ω9-D-Lys. The head group interactions with F526, L443 and R439 are maintained (Figures 7a, Figure S8 and Table S5) and the tails of C18ω5-L/D-Lys are located between TM5, TM7 and TM8, interacting with the non-polar residues (e.g., V214, L436, Y550, L557, and F567) and Y550 interacts with the lipid tail above the double bond (Figure 7), giving a structural basis for their activity.

**Figure 7.**
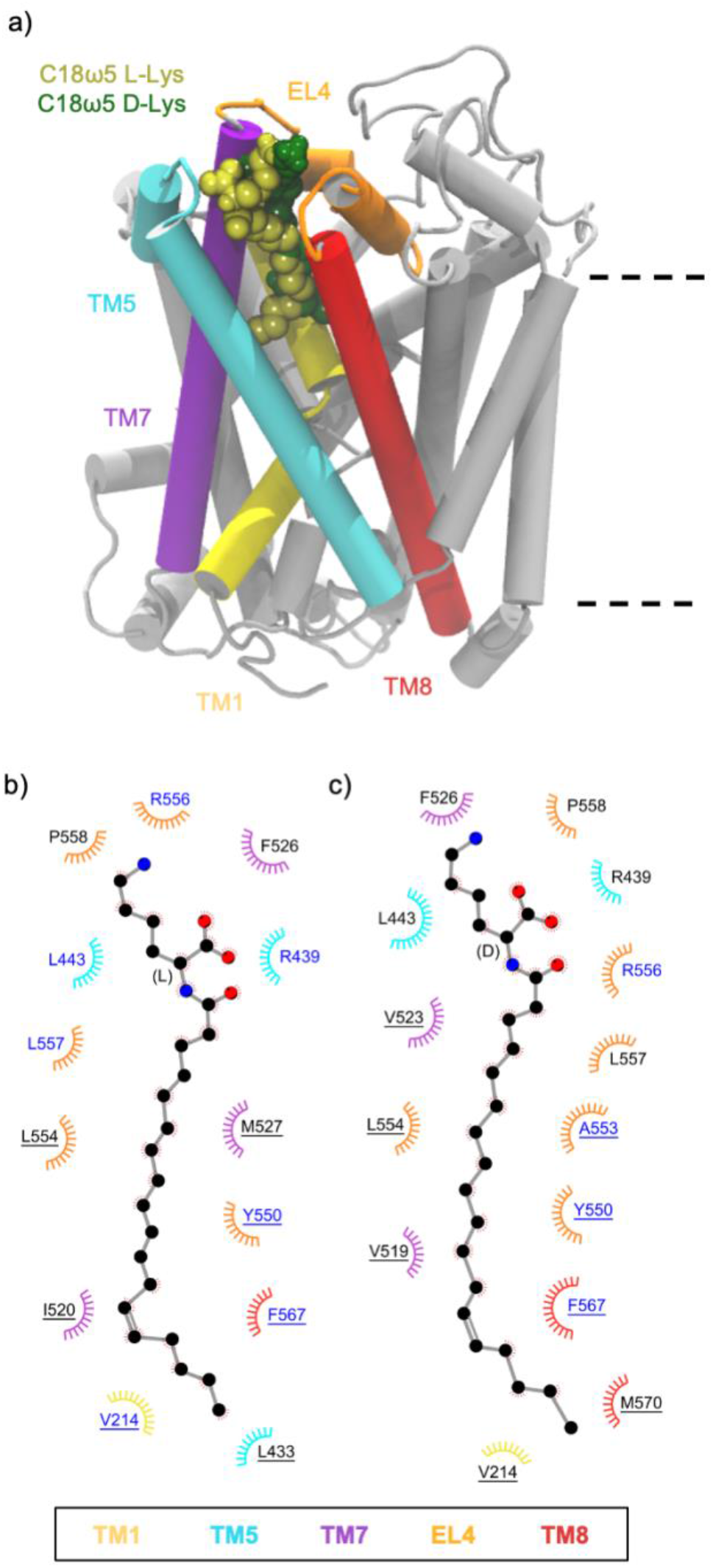
a) Positions of the C18ω9 L-Lys, C18ω5 L-Lys and C18ω5 D-Lys lipid inhibitor binding in the extracellular allosteric binding pocket of GlyT2. b and c) Residues that interact with C18ω5 L-Lys (b) or C18ω5 D-Lys (c) in representative snapshots from MD simulations of the lipid inhibitors bound in the allosteric binding site. Residues that interact with the lipid inhibitor for >75% or <75% of the total simulation time are highlighted in blue or black, respectively. Residues are split based on whether the amino acid interacts with the lipid inhibitor head-group (no underline) or tail-group (underlined).

Major differences in the binding interactions of the C16 isomers were observed, consistent with their IC_50_ concentrations. C16ω3-L-Lys, the most potent inhibitor of GlyT2 in this lipid series, remains bound in the allosteric pocket throughout the combined MD simulations, while C16ω3-D-Lys only remains bound in the extracellular allosteric binding pocket for ~1/3 of the total simulation time. In the case of C16ω3-L-Lys, the head group interacts with F526, L443 and R439, as observed for C18ω9-L-Lys (Figure 8 and Figure S9). Unlike other inhibitory lipids, the C16ω3-L-Lys tail does not adopt an extended conformation, but instead has a curled conformation in the allosteric binding site between TM5, TM7 and TM8. The curled C16ω3-L-Lys tail interacts with W215, Y550, L557, F567, and L436 (Figure 8b and Table S5). The interaction between C16ω3-L-Lys and W215 is unique and notable because W215 is directly adjacent to the glycine binding site and physically separates the extracellular allosteric site and the vestibular substrate binding site. Interactions with W215 may reflect communication between the extracellular allosteric binding site and the vestibular substrate binding site. This altered orientation and the interaction with W215 may in part explain why C16ω3-L-Lys is the most potent lipid inhibitor identified to date. In contrast, the potency of C16ω3-D-Lys is 65-fold lower (IC_50_ 699 nm) than C16ω3-L-Lys. Simulations of C16ω3-D-Lys in the extracellular allosteric binding site show the head group interacts with the membrane, as well as key amino acids (i.e., R439 and F526). The tail does not bind stably in the allosteric binding pocket between TM5, TM7 and TM8, but instead is oriented towards EL4 interacting with L557 and L436 and it readily dissociates from the binding site.

**Figure 8.**
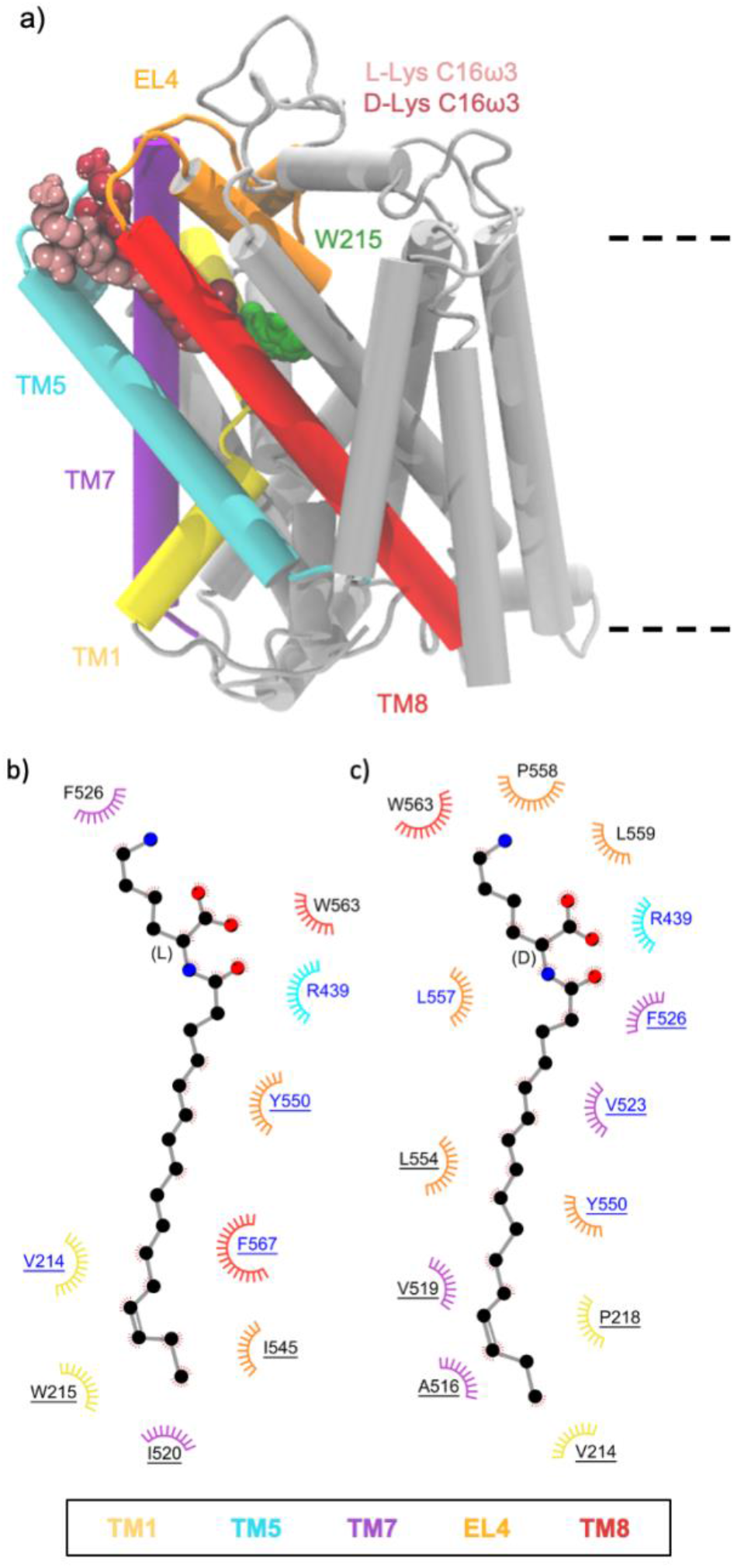
a) Positions of the C16ω7 L-Lys, C16ω3 L-Lys and C16ω3 D-Lys lipid inhibitor binding in the extracellular allosteric binding pocket of GlyT2. b and c) Residues that interact with C16ω3 L-Lys 7 (b) or C16ω3 D-Lys (c) in representative snapshots from MD simulations of the lipid inhibitors bound in the allosteric binding site. Residues that interact with the lipid inhibitor for >75% or <75% of the total simulation time are highlighted in blue or black, respectively. Residues are split based on whether the amino acid interacts with the lipid inhibitor head-group (no underline) or tail-group (underlined).

### Non-polar residues stabilize lipid inhibitor binding, while the stabilizing or destabilizing effect of cationic residues depends on the lipid inhibitor head group

The binding affinity of lipid inhibitors to the allosteric binding pocket of GlyT2 is influenced by both the amino acid head group and the depth of penetration of the lipid tail. To better understand the overall energetics of lipid inhibitor binding, the contribution of each amino acid to the relative binding energy was calculated using the MM-PBSA energy decomposition scheme. The focus was placed on binding of the most potent lipid inhibitor, C16ω3-L-Lys. Twenty-three residues were identified as having a contribution of more than −1 kJ/mol to the binding of to C16ω3-L-Lys. Twenty of these were non-polar and 3 were polar (Thr442, Glu536 and Glu459) (Figure 9).

**Figure 9.**
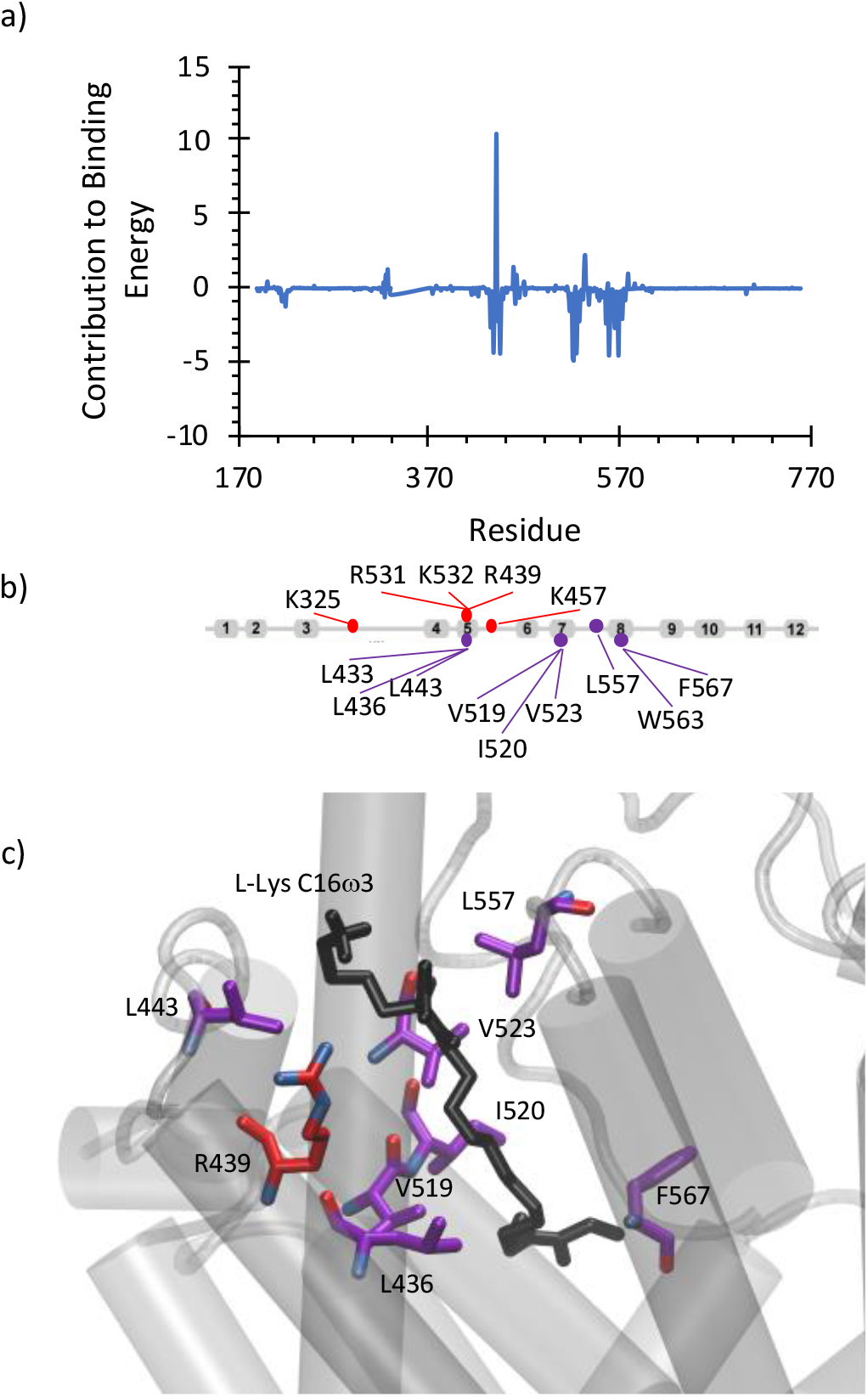
a) Energetic contribution (MM-PBSA, kJ/mol) of GlyT2 residues to C16ω3 L-Lys binding. b) Residues that destabilize the complex by > 1.2 kJ/mol or residues that stabilize the binding by > 2.5 kJ/mol and c) position of residues that destabilize the complex by > 5.0 kJ/mol or residues that stabilize the binding by > 5.0 kJ/mol relative to the C16ω3 L-Lys lipid inhibitor.

Residues that have the greatest energetic impact on binding stability are I520 (–4.9 kJ/mol), V519 (–4.7 kJ/mol), F567 (–4.5 kJ/mol), L557 (–4.5 kJ/mol), L443 (–4.4 kJ/mol), V523 (–4.4 kJ/mol) and L436 (–4.3 kJ/mol). With the exception of V519, these residues are all within 4 Å of C16ω3-L-Lys for >80% of the total simulation time (Table S5) and are mostly located around the lipid tail (Figure 9c). All residues that destabilize binding by more than 1.2 kJ/mol are cationic, while all other residues that destabilize binding by 0.1 kJ/mol are charged. R439 is the residue that most significantly destabilizes binding (10.5 kJ/mol) due to charge repulsion between the C16ω3-L-Lys head group and the R439 sidechain.

To give insight into the effect of the inhibitor structure on the activity, the overall energetics of C18ω9-L-Lys and C18ω9-L-Trp binding were also calculated (Figure 10). The two most potent inhibitors C18ω9-L-Lys and C16ω3-L-Lys have very similar binding energy decompositions to, with non-polar residues (V523, L436, I520, F526 and L557) stabilizing binding and charged amino acid destabilizing binding (R439). Head group substitution to C18ω9-L-Trp dramatically alters the per residue decomposition of the binding energy. All residues that strongly stabilize C18ω9-L-Trp binding are cationic (R556, R531, K532, K325, R439, K323 and K321) and all strongly destabilizing residues are anionic (E372, D329, E552, D322 and E530). All of these residues are positioned on the extracellular surface of protein surrounding the C18ω9-L-Trp head group. Interesting, the overall binding energy for C18ω9-L-Trp (–218.89±25.8 kJ/mol) is greater than that for the acyl lysine inhibitors (–104.5±21.5 and –110.9±29.5 kJ/mol for C16ω3-L-Lys and C18ω9-L-Lys, respectively). This result highlights that while both acyl lysine and acyl tryptophan inhibitors can act on GlyT2, the biochemical basis of inhibitor stabilization in the extracellular allosteric binding site differs between compounds.

**Figure 10.**
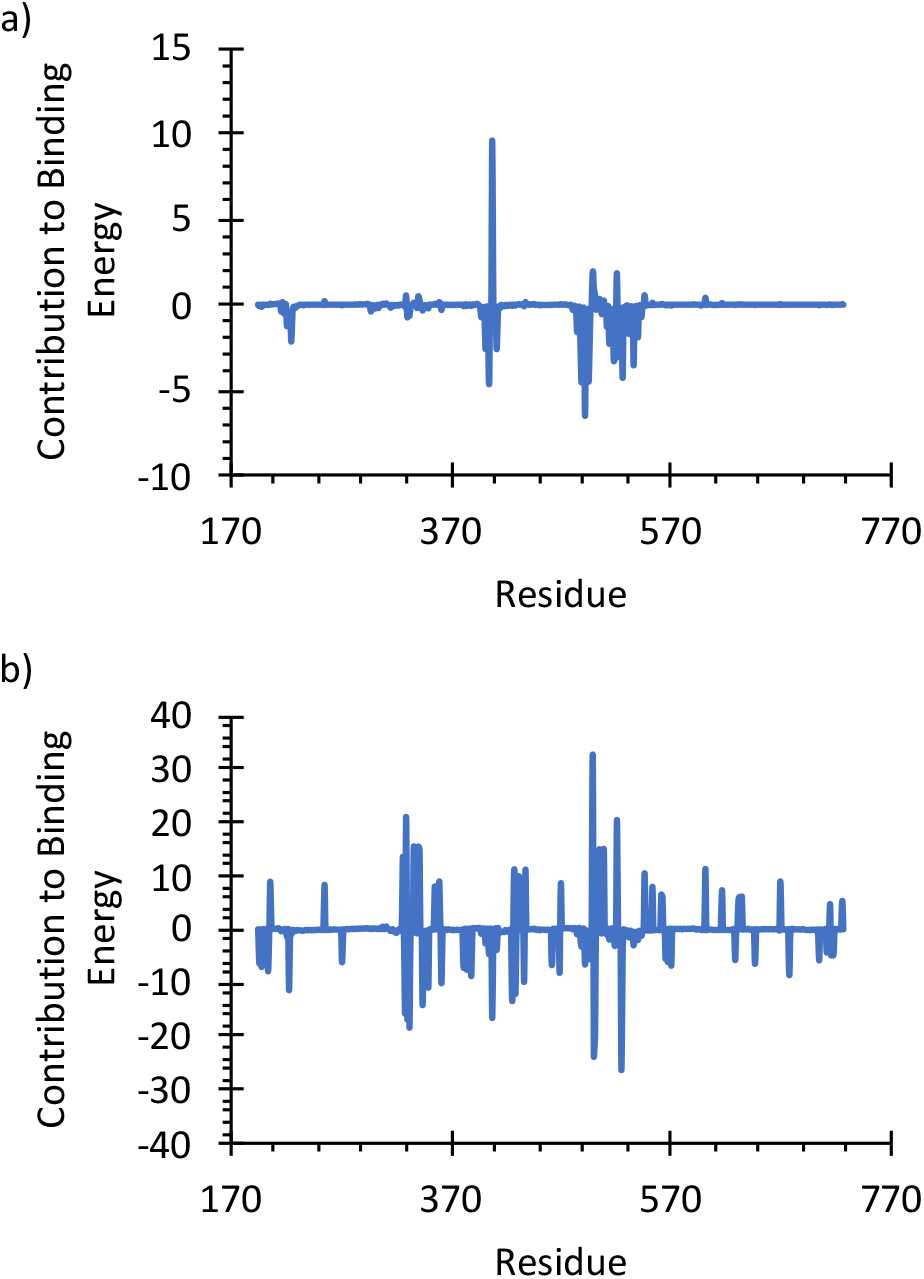
Energetic contribution (MM-PBSA, kJ/mol) of GlyT2 residues to a) OLLys and b) OLTrp binding.

## Discussion

The current work has highlighted that whilst interactions between GlyT2 and the lipid head group occur, it is the interactions with the lipid tail that are critical for determining the activity of the lipid inhibitor. This may form the basis for the potential mechanism of inhibition. The high sequence conservation of the extracellular allosteric binding site residues across SLC6 transporters suggests this lipid binding interaction could be exploited for broader inhibitor design. The position of the tail in between TM5, TM7, TM8, and EL4 spans both the core domain of the transporter (TM1, TM2, TM6, TM6 and TM7) and the scaffold domain (TM3, TM4, TM8 and TM9) (21,22) and may disrupt the rocking bundle mechanism of glycine transport. Indeed, the transition between the outward-occluded state and inward-open state in the homologous protein LeuT, has been shown to involve substantial conformational change causing the bending of TM5 and TM7 and the closure of the central vestibule by EL4 (21,22). MD simulations of the dopamine transporter and serotonin transporter have also identified changes in TM5 as a first step in this conformational transition (23). To further explore the impact the inhibitors are having on the overall protein structure, the hydrogen bonds between each of the transmembrane helices over the simulations were recorded. While most interhelical interactions remain undisrupted (i.e., TM6-TM2, TM2-TM7, TM3-TM10, TM10-TM11), notable differences are observed in the interhelical interactions when active lipid inhibitors are bound (Table S5). In the extracellular allosteric binding pocket TM5 and TM7 interact through a hydrogen bond between Y430(OH) and T512(Oγ). This interaction is lost when the active lipid inhibitors are bound but is undisturbed relative to the control when the inactive C18ω9-D-Trp is bound. This change in hydrogen bonding between TM5 and TM7 may be able to modulate the conformation change require for substrate transport. The conformations of the extracellular loops, and thereby closure of the central vestibule were also examined. The binding of the lipid inhibitors changes the conformation of EL4, which in turn changes how EL4 interacts with the other extracellular loops. While the interaction between EL4 and EL2, and EL2 and EL6 are preserved, the interaction between EL4 and EL6 is disrupted when active lipid inhibitors are bound (Figure S10). This in turn leads to the loss of a hydrogen bond in the vicinity of EL6 on the extracellular edges of TM10 (Y627 or Q630) and TM12 (Y710) and a widening of the gap between EL4 and EL6 which opens up the central vestibule. This is the first evidence of allosteric conformational changes induced by lipid inhibitor binding to GlyT2.

In the present work we have investigated the impact of head group stereochemistry and acyl tail structure on the activity of lipid-based inhibitors of GlyT2. Overall, the lipid inhibitor head-groups remain stabilised in the aromatic cage formed by R439, F526, W563 and R556, while the tail group penetrated into the cavity to different depths, dependent on inhibitor tail length and double bond position. Importantly, longer tail lengths and deeper binding correlated with increased activity. The most potent inhibitor, C16ω3-L-Lys, has unique interactions with W215, which is positioned between the extracellular allosteric binding site and substrate binding site. C16ω3-L-Lys has a potency that is similar to that of the most potent small molecule GlyT2 inhibitor to date (ORG25543, IC_50_=16 nM). The order of potency of the remaining inhibitors is C18ω9-L-Lys > C18ω9-L-Trp, C18ω9-D-Lys, C16ω7-L-Lys, C18ω5-L-Lys, C18ω5-D-Lys > C16ω7-D-Lys, C16ω3-D-Lys, C14ω5-L-Lys > C14ω5-D-Lys >>> C18ω9-D-Trp. We have shown that the energetic stabilization of acyl lysine and acyl tryptophan inhibitors in the extracellular allosteric binding site varies significantly, that overall the L-stereoisomer is typically more potent than the D-stereoisomer, and the position of the double bond influences the activity of the inhibitor. We propose that the formation of this deep binding pocket is critical for inhibition, and that bioactive lipids that can penetrate the aliphatic cavity of the extracellular allosteric binding site are superior inhibitors. Furthermore, this provides insight into the mechanism of inhibition of GlyT2 and highlights how lipids can modulate the activity of membrane proteins by infiltrating pockets between helices, a phenomenon which has been similarly proposed in cannabinoid receptors and cys-loop receptors (24–27). The current work has provided insight into the structural features of GlyT2 inhibitors that will be important for the future design of novel inhibitors of GlyT2. The structural basis of inhibitor activity reported here may provide insights that are applicable to the broader development of new compounds that could target homologous neurotransmitter transporters such as the dopamine, serotonin and noradrenaline transporter.

## Supporting Information

PDF File: Full methods, tables of average membrane properties, percentage of the total simulation time GlyT2 residues were in contact with inhibitors, positioning of bioactive lipids from simulations and hydrogen bonding interactions, per residue energy contributions for OLLys and OLTrp binding.

## Acknowledgements

This work was supported by a grant from the National Health and Medical Research Council (APP1144429). This research was undertaken with the assistance of resources and services from the National Computational Infrastructure (NCI), which is supported by the Australian Government; and the Pawsey Supercomputing Centre which is supported by the Australian Government and the Government of Western Australia.

## References

1. Goldberg, D. S., and McGee, S. J. (2011) Pain as a global public health priority. BMC Public Health 11, 770

2. Volkow, N., Benveniste, H., and McLellan, A. T. (2018) Use and misuse of opioids in chronic pain. Annu. Rev. Med. 69, 451–465

3. Todd, A. J. (2010) Neuronal circuitry for pain processing in the dorsal horn. Nature Reviews Neuroscience 11, 823–836

4. Eulenburg, V., Armsen, W., Betz, H., and Gomeza, J. (2005) Glycine transporters: essential regulators of neurotransmission. Trends in Biochemical Sciences 30, 325–333

5. Cioffi, C. L. (2018) Modulation of Glycine-Mediated Spinal Neurotransmission for the Treatment of Chronic Pain. J. Med. Chem. 61, 2652–2679

6. Mostyn, S. N., Wilson, K. A., Schumann-Gillett, A., Frangos, Z. J., Shimmon, S., Rawling, T., Ryan, R. M., O’Mara, M. L., and Vandenberg, R. J. (2019) Identification of an allosteric binding site on the human glycine transporter, GlyT2, for bioactive lipid analgesics. Elife 8, e47150

7. Caulfield, W. L., Collie, I. T., Dickins, R. S., Epemolu, O., McGuire, R., Hill, D. R., McVey, G., Morphy, J. R., Rankovic, Z., and Sundaram, H. (2001) The First Potent and Selective Inhibitors of the Glycine Transporter Type 2. J. Med. Chem. 44, 2679–2682

8. Takahashi, E., Arai, T., Akahira, M., Nakajima, M., Nishimura, K., Omori, Y., Kumagai, H., Suzuki, T., and Hayashi, R. (2014) The discovery of potent glycine transporter type-2 inhibitors: Design and synthesis of phenoxymethylbenzamide derivatives. Bioorganic & Medicinal Chemistry Letters 24, 4603–4606

9. Vandenberg, R. J., Ryan, R. M., Carland, J. E., Imlach, W. L., and Christie, M. J. (2014) Glycine transport inhibitors for the treatment of pain. Trends Pharmacol. Sci. 35, 423–430

10. Xu, T.-X., Gong, N., and Xu, T.-L. (2005) Inhibitors of GlyT1 and GlyT2 differentially modulate inhibitory transmission. Neuroreport 16

11. Mostyn, S. N., Sarker, S., Muthuraman, P., Raja, A., Shimmon, S., Rawling, T., Cioffi, C. L., and Vandenberg, R. J. (2020) Photoswitchable ORG25543 Congener Enables Optical Control of Glycine Transporter 2. ACS Chemical Neuroscience 11, 1250–1258

12. Mostyn, S. N., Rawling, T., Mohammadi, S., Shimmon, S., Frangos, Z. J., Sarker, S., Yousuf, A., Vetter, I., Ryan, R. M., Christie, M. J., and Vandenberg, R. J. (2019) Development of an N-Acyl Amino Acid That Selectively Inhibits the Glycine Transporter 2 To Produce Analgesia in a Rat Model of Chronic Pain. J. Med. Chem. 62, 2466–2484

13. Vandenberg, R. J., Mostyn, S. N., Carland, J. E., and Ryan, R. M. (2016) Glycine transporter2 inhibitors: Getting the balance right. Neurochem. Int. 98, 89–93

14. Trott, O., and Olson, A. J. (2010) AutoDock Vina: improving the speed and accuracy of docking with a new scoring function, efficient optimization, and multithreading. J. Comput. Chem. 31, 455–461

15. Stroet, M., Caron, B., Visscher, K. M., Geerke, D. P., Malde, A. K., and Mark, A. E. (2018) Automated Topology Builder Version 3.0: Prediction of Solvation Free Enthalpies in Water and Hexane. J. Chem. Theory Comput. 14, 5834–5845

16. Van Der Spoel, D., Lindahl, E., Hess, B., Groenhof, G., Mark, A. E., and Berendsen, H. J. (2005) GROMACS: fast, flexible, and free. J. Comput. Chem. 26, 1701–1718

17. Schmid, N., Eichenberger, A. P., Choutko, A., Riniker, S., Winger, M., Mark, A. E., and van Gunsteren, W. F. (2011) Definition and testing of the GROMOS force-field versions 54A7 and 54B7. Eur. Biophys. J. 40, 843

18. Humphrey, W., Dalke, A., and Schulten, K. (1996) VMD: visual molecular dynamics. J. Mol. Graphics 14, 33–38

19. Subramanian, N., Scopelitti, A. J., Carland, J. E., Ryan, R. M., O’Mara, M. L., and Vandenberg, R. J. (2016) Identification of a 3rd Na+ Binding Site of the Glycine Transporter, GlyT2. PLOS ONE 11, e0157583

20. Carland, J. E., Thomas, M., Mostyn, S. N., Subramanian, N., O’Mara, M. L., Ryan, R. M., and Vandenberg, R. J. (2017) Molecular determinants for substrate interactions with the glycine transporter GlyT2. ACS chemical neuroscience 9, 603–614

21. Forrest, L. R., Zhang, Y.-W., Jacobs, M. T., Gesmonde, J., Xie, L., Honig, B. H., and Rudnick, G. (2008) Mechanism for alternating access in neurotransmitter transporters. Proc. Natl. Acad. Sci. U. S. A. 105, 10338–10343

22. Krishnamurthy, H., and Gouaux, E. (2012) X-ray structures of LeuT in substrate-free outward-open and apo inward-open states. Nature 481, 469–474

23. Zeppelin, T., Ladefoged, L. K., Sinning, S., Periole, X., and Schiøtt, B. (2018) A direct interaction of cholesterol with the dopamine transporter prevents its out-to-inward transition. PLoS Comp. Biol. 14, e1005907

24. Hurst, D. P., Grossfield, A., Lynch, D. L., Feller, S., Romo, T. D., Gawrisch, K., Pitman, M. C., and Reggio, P. H. (2010) A lipid pathway for ligand binding is necessary for a cannabinoid G protein-coupled receptor. J. Biol. Chem. 285, 17954–17964

25. Hua, T., Vemuri, K., Pu, M., Qu, L., Han, G. W., Wu, Y., Zhao, S., Shui, W., Li, S., Korde, A., Laprairie, R. B., Stahl, E. L., Ho, J. H., Zvonok, N., Zhou, H., Kufareva, I., Wu, B., Zhao, Q., Hanson, M. A., Bohn, L. M., Makriyannis, A., Stevens, R. C., and Liu, Z. J. (2016) Crystal Structure of the Human Cannabinoid Receptor CB(1). Cell 167, 750–762.e714

26. Hanson, M. A., Roth, C. B., Jo, E., Griffith, M. T., Scott, F. L., Reinhart, G., Desale, H., Clemons, B., Cahalan, S. M., Schuerer, S. C., Sanna, M. G., Han, G. W., Kuhn, P., Rosen, H., and Stevens, R. C. (2012) Crystal structure of a lipid G protein-coupled receptor. Science 335, 851–855

27. Bai, J. Y., Ding, W. G., Kojima, A., Seto, T., and Matsuura, H. (2015) Putative binding sites for arachidonic acid on the human cardiac Kv 1.5 channel. Br J Pharmacol 172, 5281–5292.

